# Combinatorial microRNA loading into extracellular vesicles for anti-inflammatory therapy

**DOI:** 10.1101/2022.07.13.499941

**Authors:** Alex Eli Pottash, Daniel Levy, Anjana Jeyaram, Leo Kuo, Stephanie M. Kronstadt, Wei Chao, Steven M. Jay

## Abstract

Extracellular vesicles (EVs) have emerged as promising therapeutic entities in part due to their potential to regulate multiple signaling pathways in target cells. This potential is derived from the broad array of constituent and/or cargo molecules associated with EVs. Among these, microRNAs (miRNAs) are commonly implicated as important and have been associated with a wide variety of EV-induced biological phenomena. While controlled loading of single miRNAs is a well-documented approach for enhancing EV bioactivity, loading of multiple miRNAs has not been fully leveraged to maximize the potential of EV-based therapies. Here, an established approach to extrinsic nucleic acid loading of EVs, sonication, was utilized to enable controlled loading of multiple miRNAs in HEK293T EVs. Combinations of carefully chosen miRNAs were compared to single miRNAs with respect to anti-inflammatory outcomes in assays of increasing stringency, with the combination of miR-146a, miR-155, and miR-223 found to have the most potential amongst tested groups.

## Introduction

Inflammation-related diseases are responsible for millions of deaths every year [1, 2]. While inflammation is a critical part of an effective response to harmful stimuli, inappropriate acute or chronic inflammatory signaling can cause harm to the body. Widespread adoption of inflammation management protocols has helped lower death rates, but there are still many inflammatory disorders for which there are no specific approved treatments. As a result, new therapeutic approaches are being pursued. An emerging strategy involves microRNAs (miRNAs), which have been shown to play significant roles in inflammation in general and in specific inflammatory conditions such as sepsis, both in promoting pathogenesis as well as recovery [3–15]. For example, miRNAs such as miR-146a and miR-223 have been shown to be downregulated in both septic vs. healthy patients and in non-surviving vs. surviving patients [4]. Thus, the concept of therapeutic miRNA delivery is intriguing as a possible novel anti-inflammatory treatment.

When considering potential vehicles for miRNA delivery, extracellular vesicles (EVs) have been implicated as promising based on their reported natural ability to facilitate intercellular RNA transfer [16]. While the physiological significance of EV-mediated miRNA transfer is still controversial [17–19], the capabilities of specifically-loaded EVs for small RNA delivery (siRNA and miRNA) have been clearly established [20–22]. Further, direct comparisons of EVs and other potential miRNA delivery vehicles such as liposomes have indicated the potential superiority of EVs [23–25]. Thus, EV-mediated miRNA delivery to treat inflammation is worthy of focused investigation.

Here, we built on previous work from our group using a sonication-based miRNA loading strategy to package miRNA into EVs without requiring chemical modifications [17]. Our prior study, like many in the field to date, investigated delivery of only a single miRNA species. In this work, we sought to exploit the potential synergy of regulating multiple anti-inflammatory pathways by loading multiple miRNA species into a single EV population. Combinations of miRNAs were tested in an *in vitro* macrophage inflammation model, which was previously shown to correlate with *in vivo* outcomes for EVs [26]. Finally, the most effective combination was tested in an *in vivo* endotoxemia model.

## Materials and Methods

### Cell culture

Human embryonic kidney HEK293T cells and RAW264.7 mouse macrophage cells were cultured in Dulbecco’s modified Eagle’s medium (DMEM) (Corning 10-013-CV; Corning, NY, USA) supplemented with 10% EV-depleted fetal bovine serum (FBS) and 1% penicillin/streptomycin in T175 tissue culture polystyrene flasks. FBS was EV-depleted via 100,000 x g centrifugation at 4C for 16h where the supernatant was retained.

### Extracellular vesicle isolation

Conditioned media was collected and subjected to differential centrifugation. Briefly, the supernatant was centrifuged at 1,000 x g for 10min, 2,000 x g for 20min, 10,000 x g for 30min, for each of which the supernatant was retained, and finally, 100,000 x g for 2h, after which the pellet was resuspended in PBS and collected. The final spin was performed using a Beckman Optima L-90K ultracentrifuge with T70i rotor. This resuspension was washed 2x using Nanosep 300-kDa MWCO spin columns (OD300C35; Pall, Port Washington, NY, USA). The washed EVs were resuspended in PBS and filtered using an 0.2-μm syringe filter. EV size distribution and concentration were determined by nanoparticle tracking analysis (NTA) via a NanoSight LM10. Each sample was analyzed in triplicate using consistent acquisition settings. Total EV protein was determined via bicinchoninic acid assay (BCA) following manufacturer’s protocol. Relative levels of relevant protein components were determined via western blotting. Alix (ab186429; Abcam), TSG101 (ab125011; Abcam), GAPDH (2118L; Cell Signaling Technology) and CD63 (25682-1-AP; Thermo Fisher). Primary antibodies were added at a 1:1000 dilution, except for GAPDH (1:2000). Secondary antibody IRDye 800CW anti-Rabbit (926-32211; LI-COR Biosciences, Lincoln, NE, USA) were added at 1:10000 dilution, and membranes were imaged on a LI-COR Odyssey CLX Imager.

### Extracellular vesicle loading

100μg EVs, corresponding to ∼3e9 particles detected by NTA, were mixed with 1nmol miRNA mimic and the volume was brought up to 100μl with PBS. This mixture was incubated for 30m at room temperature, before being sonicated at in a water bath sonicator (VWR® symphony™; cat# 97043-964, 2.8L capacity, dimensions 24L × 14W × 10D cm) at 35kHz for 15s, placed on ice for 1m, and sonicated for a second 15s. The mixture was placed back on ice briefly, then washed 3x using Nanosep 300-kDa MWCO spin columns to remove unincorporated RNA and resuspended by PBS. The miRNA mimics (Dharmacon) used were: hsa-miR146a-5p (C-300630-03); hsa-miR-155-5p (C-310430-07); hsa-miR-223-3p (C-300580-07); Negative Control #1 (C-310391-05).

### Transmission electron microscopy (TEM)

EVs were negatively stained using a protocol as previously described [318]. Briefly, 4% paraformaldehyde (10μL) was added to EVs (10μL) and incubated for 30min. A carbon film grid was placed on the paraformaldehyde/EV droplet for 20min, and washed with PBS. Then, the grid was placed on 1% glutaraldehyde (50μL) for 5min, and washed eight times with water. Finally, the grid was placed on uranyl acetate replacement stain (50μL) for 10min, and left to dry for 10min.

### Fluorescently-labeled RNA co-loading test

Cy3-labeled miR-93 (CTM-433488) and Cy5-labeled miR-126 (CTM-508110) (Dharmacon) were loaded at indicated ratios according to the normal sonication protocol. After washing, fluorescence readings were taken and normalized to total fluorescence.

### *In vitro* RAW264.7 inflammatory assay

RAW264.7 cells were seeded in DMEM supplemented with 5% FBS in a 48-well plate at 100,000 cells per well. All EVs were prepared by sonication and doses were normalized by protein content after sonication and washing. All treatments were diluted in DMEM supplemented with 5% FBS. In the “Pre-treat” regime, cells were treated with EVs or PBS alone for 24 hours, when supernatant was replaced by media with 10ng/mL LPS for 4h. In the “Co-treat” regime, both EV treatments and 10ng/mL LPS were added concomitantly for 24 hours. In the “Post-treat” regime, cells were treated with 10ng/mL LPS for 24 hours, and then 10ng/mL LPS with EV treatments for another 24 hours. After all final treatments, media was collected and stored at −80C. IL-6 concentration was determined using the Mouse IL-6 DuoSet ELISA Kit (R&D Systems; DY406). Cell transfection was achieved using HiPerfect (Qiagen) according to manufacturer’s protocol. For the “Co-treat” regime, phagocytosis was measured after the removal of the media, following the Vybrant Phagocytosis Assay Kit (Invitrogen) manufacturer’s protocol. All tests were performed in biological triplicate.

### ELISA

Samples were diluted as needed and cytokine concentrations were determined via DuoSet ELISA Kits (R&D Systems): IL-6 (DY406), TNFa (DY410), MIP-2 (DY452), IL-1β (DY401), and CCL22 (DY439).

### Proteome array

An antibody-based protein array was performed on cell supernatants after a “Pre-treat” regime, using Proteome Profiler Mouse XL Cytokine Array (R&D Systems) according to manufacturer’s protocol. Expression was normalized between membranes.

### *In vivo* endotoxemia study

Male C57BL/6J mice (Jackson Labs), 8 to 12 weeks of age, were used in this study. The animals were kept at a constant temperature (25°C) under a 12h light/dark cycle with free access to food and water. On the first and second day, animals received a 200μl intraperitoneal injection of PBS or sonicated EVs at a concentration of 2.1e10 particles/mL (by NTA). On the third day, animals received an intraperitoneal injection of 5mg/kg LPS. Three hours later, animals were anesthetized and sacrificed via cardiac blood collection. Blood was collected into EDTA-coated tubes (450480; Greiner Bio-One) and spun at 1,000 x g for 15min to produce plasma. All animal work was carried out in accordance with the NIH guidelines and approved by the Institutional Animal Care and Use Committee (IACUC) at the University of Maryland College Park.

### Statistical analysis

Data are presented as mean ± SD. One-way ANOVAs with Dunnett’s multiple comparison test to determine statistical significance in *in vitro* inflammatory assay, and *in vivo* endotoxemia experiments. All statistical analysis was performed with Prism 8 (GraphPad Software, La Jolla, CA).

## Results

### EV loading and characterization

To test multiple different combinations of miRNAs, a method of sonication-mediated EV loading previously developed by our lab was employed [27]. The sonication method is an exogenous loading technique in which pre-synthesized siRNA or miRNA mimics can be mixed in any combination to EVs and loaded, with minimal damage to both the EVs and RNA [27]. EVs derived from HEK293T cells were collected and the ability to controllably co-load two different miRNA cargos into a single EV population was determined by mixing and sonicating Cy3-labeled miR-93 and Cy5-labeled miR-126 in varying proportions (Fig. 1A). These two RNAs with differing sequences were found to be loaded near their input proportion. Sonicated EVs were characterized via western blot (Fig. 1B), nanoparticle tracking analysis (NTA) (Fig. 1C), and transmission electron microscopy (TEM) (Fig. 1D) according to the recommendations of the International Society for Extracellular Vesicles [28].

**Figure 1.**
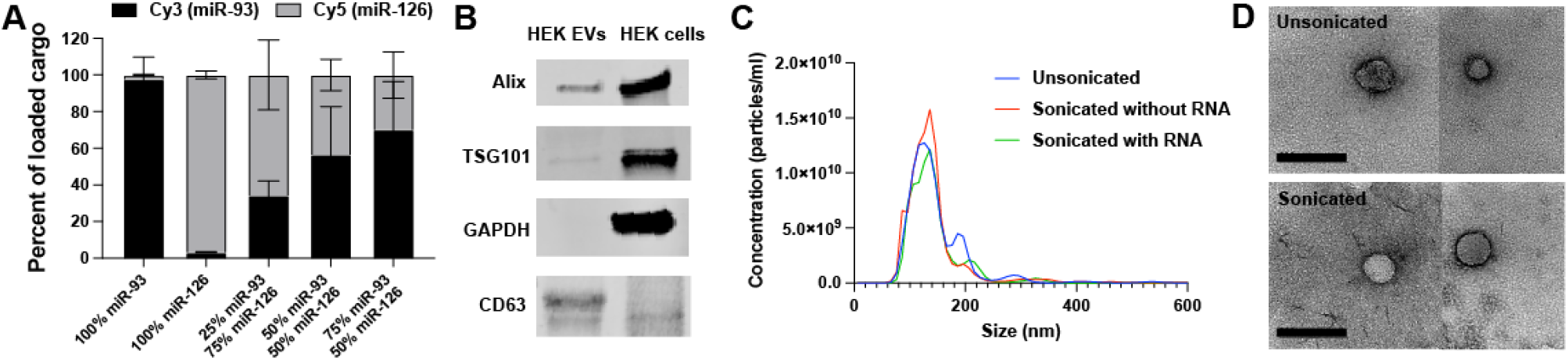
EV characterization and co-loading validation. (A) Relative quantification of co-loaded fluorescently-tagged miRNA mimics. (B). Western blot of EVs vs parental cells. (C) Nanoparticle Tracking Analysis (NTA) performed on EVs unsonicated, sonicated, and sonicated with miRNA. (D) Transmission electron micrographs (TEM) of unsonicated and sonicated EVs. Scale bar = 200nm.

### Screening for anti-inflammatory miRNA

As an initial assessment of anti-inflammatory bioactivity, effect on IL-6 secretion was selected as a screening criterion based on a prior report that showed correlation between the effects of EVs on IL-6 secretion in vitro and their anti-inflammatory activity *in vivo* [26]. HEK293T EVs were chosen for study due to their expected limited anti-inflammatory bioactivity as well as low intrinsic RNA content [29]. EVs were loaded with three different miRNA mimics (miR-146a, miR-155, and miR-223; Dharmacon) that were found in literature to be downregulated in septic patients, to regulate the TLR4 inflammatory pathway, and/or to have altered expression levels in response to LPS stimulation [4, 30–36]. While certain single-stranded miRNAs (including miR-146a-5p) have been shown to be pro-inflammatory TLR agonists [3, 9, 37, 38], these double-stranded mimics are designed to interact with the RNA-induced silencing complex (RISC) with preferential strand selection. The miRNAs were loaded either individually, in combination with another, or as the complete group. In this way, each miRNA could be compared with others both as a mono-treatment and when left out of the complete group. These EV treatments were applied to RAW264.7 murine macrophage cells for 24 hours, when supernatant was replaced by LPS treatment for 4 hours, in a “Pre-treat” regime. At the end of the LPS treatment, supernatants were collected, assessed using an IL-6 ELISA, and compared to the “No miRNA” group. All treatment groups led to significant anti-inflammatory effects, with dose-dependence evident (Fig. 2A). Interestingly, the No miRNA group (unmodified HEK293T EVs) showed an anti-inflammatory effect on par with Dex, reflecting prior data showing benefits of HEK293T EVs in a sepsis model via an unknown mechanism [39]. Cell phagocytic behavior was tested after LPS treatment to see if EV-mediated miRNA treatment impaired phagocytosis (Fig. 2B). No significant changes were detected, indicating that treatments were not inducing endotoxin tolerance. Due to the effectiveness of each miRNA combination, more challenging regimes were employed to differentiate between combinations.

**Figure 2.**
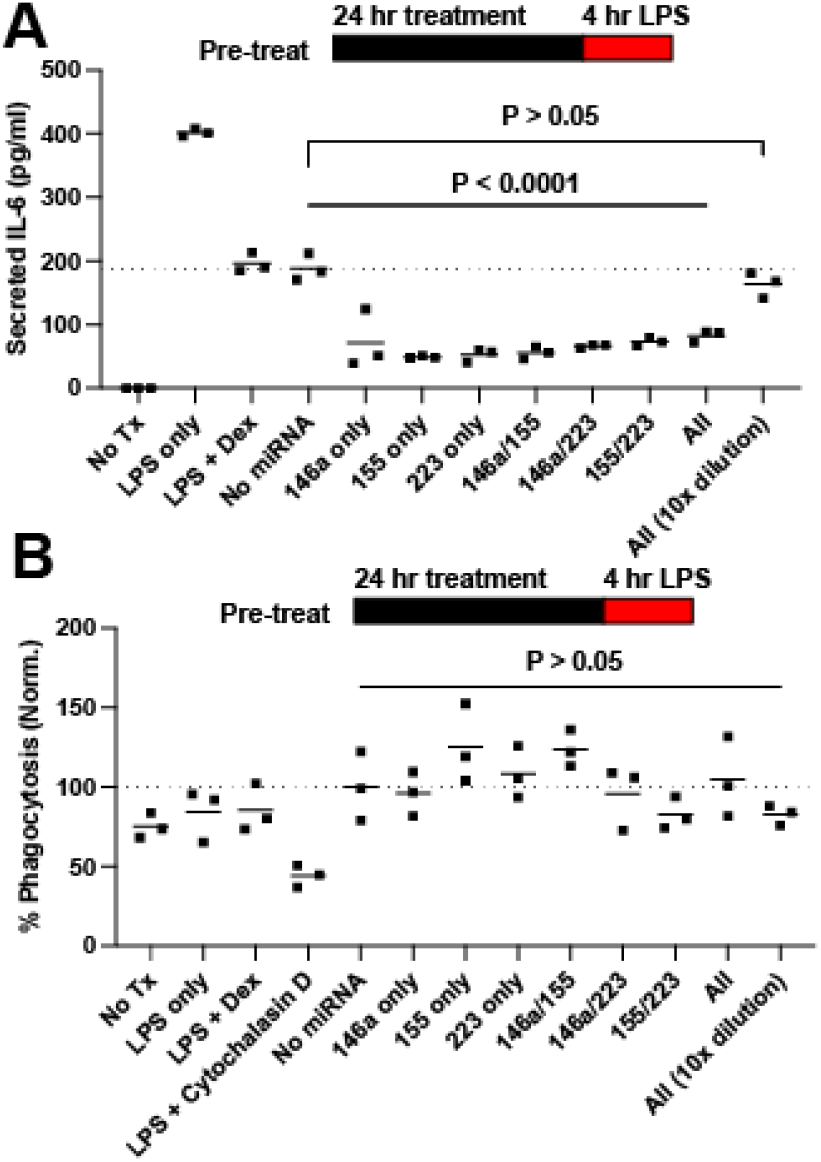
Screening of miRNA for anti-inflammatory combination. (A) Secreted IL-6 in response to LPS in a pre-treatment regime. “All” refers to all miRNA listed. (B) Phagocytosis as measured by the Vybrant Phagocytosis Assay Kit (Invitrogen).

All groups were next tested in a “Co-treat” regime, wherein LPS and EV treatments were both applied concurrently to RAW264.7 cells for 24 hours (Fig. 3A). miR-146a alone had a significant anti-inflammatory effect, while miR-223 alone and miR-155 alone had no effect. In contrast, strikingly, the 155/223 combination significantly reduced IL-6 secretion. The 146a/223 treatment was not significantly effective, while the 146a/155 and 146a/155/223 treatments significantly reduced IL-6 secretion. Next, all groups were tested in a “Post-treat” regime, wherein LPS was applied concurrently to RAW264.7 cells for 24 hours, and then LPS and EV treatments were concurrently applied for 24 hours (Fig. 3B). In this regime, no significant effects were detected except for with the complete combination of 146a/155/223. Finally, a murine endotoxemia model was employed to assess the 146a/155/223 combination. IL-6 serum levels in mice dosed with 146a/155/223 showed a 23% reduction in cytokine plasma concentration compared to LPS control animals. (P=0.07) (Fig. 3C). Interestingly, a serum IL-6 reduction associated with the control group, HEK EVs loaded with cel-miR-67 miRNA mimic (Ctrl), was also observed, once again reflecting previous data showing unexpected anti-inflammatory benefits of HEK293T EVs [39].

**Figure 3.**
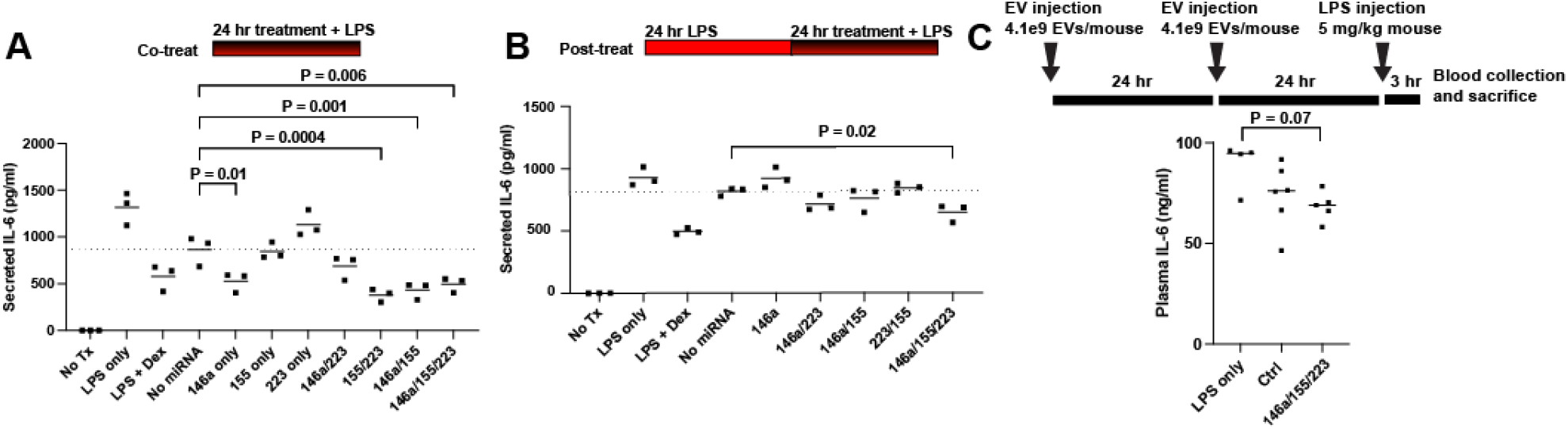
Screening of miRNA for anti-inflammatory combination. (A) Secreted IL-6 in response to LPS in a co-treatment regime. (B) Secreted IL-6 in response to LPS in a post-treatment regime. (C) IL-6 plasma levels in endotoxemic mice. Results were analyzed via one-way ANOVA.

### Delivery of miRs 146a/155/223 has variable anti-inflammatory effects aside from reducing IL-6 secretion

Given the effectiveness of the miR 146a/155/223 combination (“TRI”) in suppressing IL-6 secretion, we screened to see if other relevant secreted cytokines were also regulated using an antibody-based cytokine array. Pre-treatment of RAW264.7 cells with EV-delivered TRI or HEK EVs loaded with cel-miR-67 Negative Control miRNA mimic (“NC”) showed differential protein expression after LPS treatment for 4 hours (Fig. 4A). In comparison to NC, TRI induced downregulation of IL-6, IL-10, CCL22, CCL17, CXCL10, CXCL13 and CXCL16 (Fig. 3b). Array data for all targets is available in Spreadsheet S1. CCL22 downregulation in vitro was verified via ELISA (Fig. 4C). However, TRI treatment failed to induce any change in CCL22, TNFa, MIP-2, or IL-1β secretion in endotoxemic mice as compared to NC or LPS-only controls (Fig. 4D).

**Figure 4.**
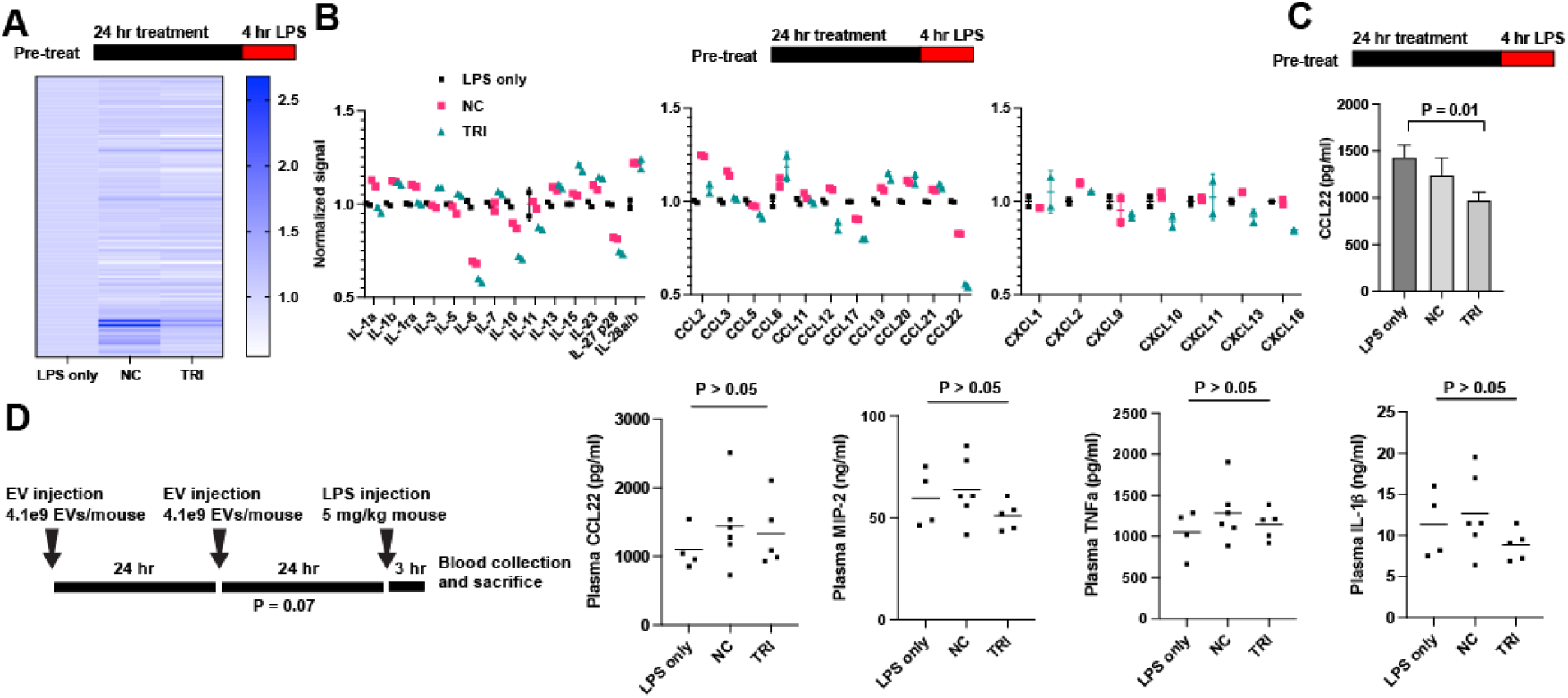
Screen for extracellular protein targets of the miR 146a/155/223 combination (“TRI”). (A) Relative expression for all targets. (B) Relative expression for IL, CCL, CXCL cytokines. Expression as measured by the Proteome Profiler Mouse XL Cytokine Array (R&D Systems). (C) CCL22 expression was quantified via ELISA. Results were analyzed via one-way ANOVA. (D) Treatment schedule and serum cytokine levels for endotoxemic mice pre-treated with TRI or NC as indicated. Results were analyzed via one-way ANOVA.

## Discussion

We had previously established [27] that sonication enables the loading of miRNA into EVs with only slight diminishment of *in vitro* EV uptake compared to unmodified EVs. In this study, we showed that the loading of multiple small RNA sequences by sonication is predictable based on the proportion of their concentration in solution. This technique may thus allow several advantages over competing EV loading strategies. Any mixture of miRNA sequences can potentially be loaded into a single EV population with a reproducible loading efficiency. As opposed to mixtures of singly-loaded EVs, the pre-mixing of miRNA allows for the possibility of loading multiple miRNAs into a single vesicle, promoting proportional delivery to a recipient cell. This exogenous loading technique is also adaptable for any small RNA cargo and does not require any manipulation of the cargo or producer cells.

To take advantage of this system, we performed a screen for anti-inflammatory miRNA combinations using a limited number of miRNAs selected from the literature. These miRNA combinations were passed through progressively more rigorous LPS challenges *in vitro* to determine if any specific combination of miRNAs is superior in reducing inflammation. That process identified the combination of miR-146a, miR-155, and miR-223 as being the most efficacious in reducing IL-6 production by RAW264.7 macrophages in response to LPS. This finding echoes work by Bhaskaran *et al*. that found that overexpression of three miRNAs in glioblastoma had a combinatorial anti-cancer effect [40], as well as a clinical study by Marik *et al*. which found that a combination of hydrocortisone, ascorbic acid, and thiamine worked synergistically as an anti-inflammatory against sepsis [41, 42].

miR-146a, miR-155, and miR-223 have been studied as anti-inflammatory miRNAs that change expression levels in response to LPS and target proteins in the TLR4 pathway [30–32, 43]. Interestingly, these miRNA targets are largely non-overlapping, perhaps indicating that when attempting to downregulate a cellular pathway, greater effect may be achieved by targeting different proteins in that pathway rather than focusing on one protein. Work by *Schulte et al*. described the tiered response by macrophages to LPS, in which miR-146 expression saturates at even sub-inflammatory LPS concentrations in order to protect against hyper-sensitivity, whereas miR-155 is expressed proportionally over a broad range of LPS concentrations in order to respond appropriately to the level of stimulation [44]. This indicates that both miRNAs seem to work in tandem to prevent an extreme cellular response. However, in other contexts, introducing miR-155 has been shown to be pro-inflammatory [45, 46]. For example, EVs from wild-type bone marrow-derived dendritic cells (BMDCs) increased IL-6 production in response to LPS in miR-155^-/-^ BMDCs and mice, compared to EVs from miR-155^-/-^ BMDCs [47]. These seemingly contradictory results indicate that miR-155 activity is nuanced and likely very context dependent. Concurrent introduction of other anti-inflammatory miRNAs like miR-146a and miR-223 may tilt the RNA network towards an environment in which miR-155 suppresses inflammation.

The results of our protein array showed a downregulation of IL-6, as expected. CCL22 and CCL17, the two CCR4 ligands, which are involved in T-cell chemotaxis, are also downregulated by the TRI treatment. Interestingly, in an LPS challenge model, CCR4^-/-^ mice had decreased cytokine release and higher survival rate when compared to wild-type mice [48]. In another study, CCR4^-/-^ mice had reduced immune response and greater survival after cecal ligation and puncture (CLP), and greater responsiveness and survival to a secondary fungal challenge [49]. These previous results indicated that, in addition to inhibition of IL-6, inhibition of the CCR4 ligands CCL22 and CCL17 may lead to an improved outcome *in vivo*.

Despite these encouraging signs, we saw no significant decrease in pro-inflammatory cytokines in response to TRI *in vivo*. There are multiple reasons why this may be the case. Firstly, since cell source plays a role in EV biodistribution and delivery [50], the choice of HEK293-derived EVs may limit an *in vivo* effect. A recent study showed that HEK293 EVs have a short half-life in healthy mice; in one hour, 80% of EVs were cleared from circulation [51]. It is possible that cargo packaged within EVs from mesenchymal stromal cells or another cell source could have a greater chance of functional delivery. Additionally, while sonication may inhibit EV delivery only slightly *in vitro*, this effect may be increased under more challenging delivery conditions i*n vivo*. Finally, the *in vitro* model used to screen for anti-inflammatory effects may be insufficiently representative of *in vivo* dynamics, despite prior correlation noted in the literature [26]. For example, the RAW264.7 macrophage model may be insufficiently representative of native macrophage behavior and is certainly insufficiently representative of other cell types affected by LPS injection.

## Conclusion

Sonication is an effective method for loading multiple miRNAs into EVs in predictable proportions. Given the vast number of targets that are regulated by any one miRNA sequence, it would be difficult to fully map or predict the changes in the transcriptome, proteome, or phenotype of a cell that takes up one miRNA, let alone three. In this way, while literature can guide selection of therapeutic miRNA, empirical combinatorial testing of multiple miRNAs may be necessary when seeking to design a miRNA-based therapeutic. This work, which by no means exhausts the possible space of miRNA combinations, is nonetheless our attempt to illuminate the strengths of such an approach.

## Supplementary Materials

**Figure S1:** Original western blot images.

**Spreadsheet S1:** Normalized protein array data, as displayed in Figure 4A.

## Author Contributions

Conceptualization, A.E.P. and S.M.J.; Methodology, A.E.P.; Validation, A.E.P.; Formal Analysis, A.E.P.; Investigation, A.E.P, D.L., A.J., L.K., and S.M.K.; Resources, S.M.J.; Data Curation, A.E.P.; Writing – Original Draft Preparation, A.E.P.; Writing – Review & Editing, S.M.J.; Visualization, A.E.P.; Supervision, W.C. and S.M.J.; Project Administration, S.M.J.; Funding Acquisition, S.M.J.

## Data Availability Statement

The normalized protein array data displayed in Figure 4A are available in Spreadsheet S1.

