## Supplementary figures and images for "Combinatorial microRNA loading into extracellular vesicles for anti-inflammatory therapy"

### Figure S1

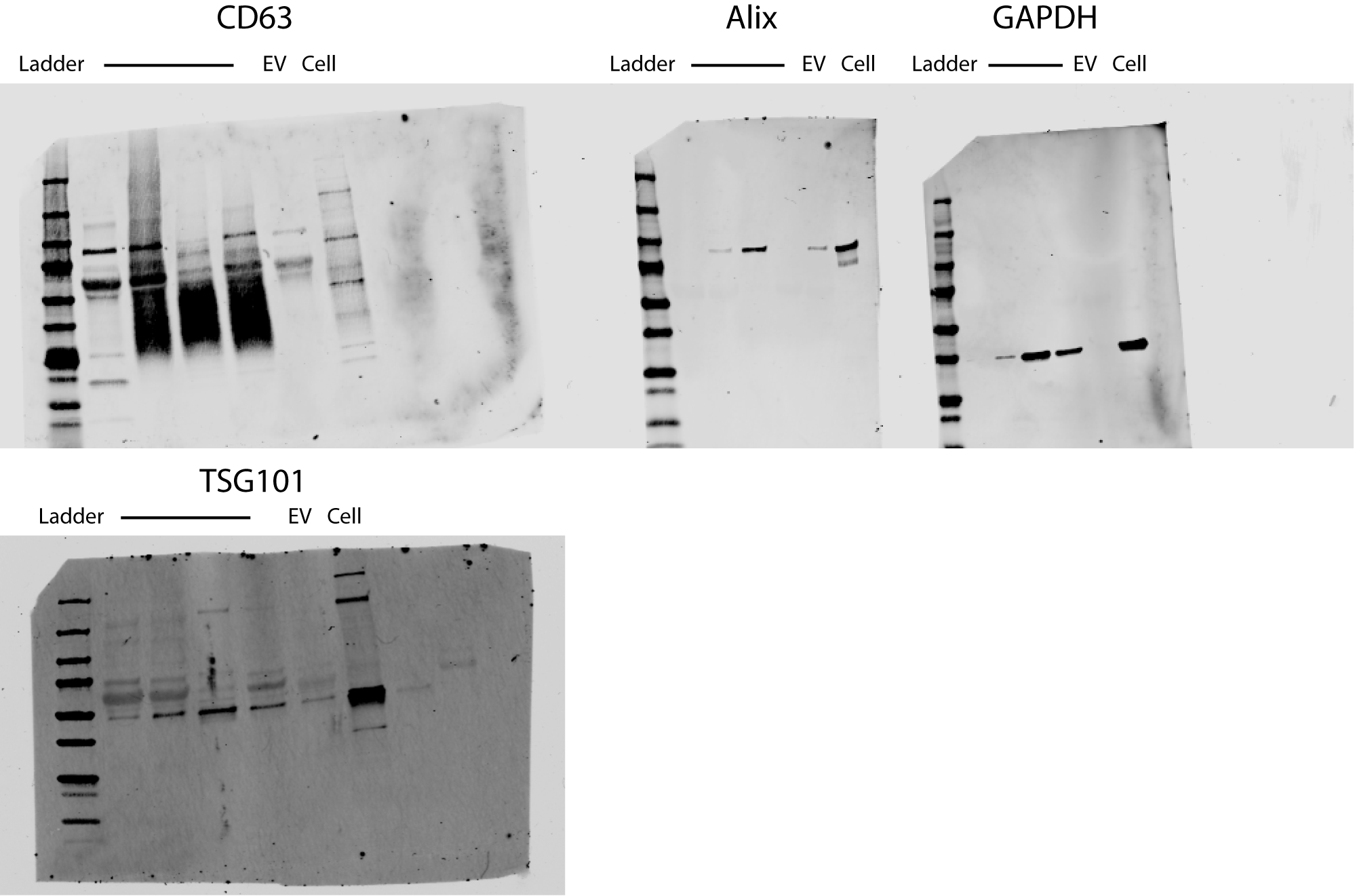
